# Predictors of protein evolution in the drosophilid immune system

**DOI:** 10.1101/2025.09.11.675540

**Authors:** Pankaj Dhakad, Darren J. Obbard

**Affiliations:** Institute of Ecology and Evolution, University of Edinburgh, Edinburgh, United Kingdom

## Abstract

The evolutionary dynamics of immune genes are shaped by diverse selective pressures, yet the relative roles of gene-level traits, functional specialization, and pathway context remain poorly understood. Here, we applied a meta-analytic mixed model approach to quantify how immune-pathway genes differ from other genes in their rates of protein sequence divergence (dN/dS), evidence for positive selection, and gene turnover rate (λ), while simultaneously accounting for gene length, expression level, genetic and protein–protein interactions, and structural features such as relative solvent accessibility (RSA). In general, rates of sequence evolution were strongly and positively associated with RSA, and negatively with gene length, expression, and genetic/protein-protein interactions, while gene turnover rate was largely unaffected by these factors. We find immune genes evolved significantly faster at the protein sequence level than non-immune genes, but contrary to our expectation exhibited lower gene turnover rates. Functional and pathway-level analyses revealed accelerated evolution in effectors, receptors, and antiviral genes, with cGAS–STING and Toll pathways showing the highest dN/dS. Gene turnover rate was elevated only in effectors, whereas cellular defence genes were particularly conserved. We also found evidence for elevated proportion of sites under episodic positive selection in immune genes, particularly in effectors, indicating ongoing adaptive diversification. These findings highlight how immune diversification in Drosophilidae arises from multiple, partly independent evolutionary axes, shaped jointly by structural constraints, functional roles, and lineage-specific pathogen pressures.

## Introduction

The evolution of immune genes is widely thought to be shaped by a persistent arms race between hosts and their pathogens. Across animals, immune-related genes are generally reported to be among the most rapidly evolving components of the genome, exhibiting elevated rates of non-synonymous substitution, gene duplication and loss, and structural innovation that often exceed genomic background levels (Nielsen, et al. 2005; Sackton, et al. 2007; Obbard, Welch, et al. 2009; Shultz and Sackton 2019; Vinkler, et al. 2023). This rapid evolution is often interpreted as a hallmark of coevolution, where pathogens exert recurrent selective pressure, driving adaptive changes in recognition, signalling, and effector components of the immune system (Sackton, et al. 2007; Sironi, et al. 2015; Świderská, et al. 2018; Velová, et al. 2018; Shultz and Sackton 2019; Lazzaro, et al. 2020; Davies, et al. 2021).

In *Drosophila melanogaster*, the innate immune system is well characterized and has served as a foundational model for understanding both the mechanisms and evolution of immunity (Lemaitre and Hoffmann 2007), especially of invertebrates. The *Drosophila* immune system is composed of both cellular and humoral immune responses to pathogens. The cellular response comprises a set of differentiated hemocyte populations that are responsible for phagocytosis of microbes by plasmatocytes, and encapsulation and melanisation of larger parasites by lamellocytes and crystal cells respectively (Honti, et al. 2014; Balog, et al. 2021)—although there is substantial variation among species (Salazar-Jaramillo, et al. 2014a; Cinege, et al. 2024). The humoral component consists of the Toll and Imd NF-κB pathways, while additional components include the JAK-STAT cytokine pathway, JNK pathway, as well as RNAi and cGAS–STING antiviral systems (reviewed in Westlake, et al. 2024). Broadly, these different pathways respond to distinct pathogen classes. Toll signalling primarily targets Gram-positive bacteria and fungi, Imd targets Gram-negative bacteria, RNAi and cGAS-STING act against viral nucleic acids, and JAK-STAT is activated by cellular stress or wounding (Gottar, et al. 2002; Wang, et al. 2006; Myllymäki and Rämet 2014; Tafesh-Edwards and Eleftherianos 2020; Cai, et al. 2022; Huang, et al. 2023). However, the role of these pathways is not absolute, with evidence of crosstalk between immune pathways and overlapping roles in pathogen defence (Tanji, et al. 2007; Nishide, et al. 2019). For example, Some AMPs (such as Drosomycin) are activated by both Toll and Imd pathways (Valanne, et al. 2010). Moreover, infections with gram-positive and gram-negative bacteria can co-activate both pathways in a synergistic manner (Tanji, et al. 2007). Beyond their classical roles, both the Toll and Imd pathways have also been implicated in antiviral immunity, with Toll signalling mediated AMPs upregulated against Drosophila X virus (DXV), and Imd mediating responses to Sindbis and Cricket Paralysis viruses (Costa, et al. 2009; Sabin, et al. 2010).

In all pathways, several genes have been found to be rapidly evolving, potentially as a consequence of host-parasite arms races (Schlenke and Begun 2003; Obbard, et al. 2006; Jiggins and Kim 2007; Sackton, et al. 2007; Obbard, Gordon, et al. 2009; Palmer, et al. 2018). Genes acting at different steps in these pathways—including extracellular recognition proteins (such as PGRPs, GNBPs), intracellular signalling molecules (such as Relish, Dorsal), and effector genes (such as *Attacin*, *Defensin*)—differ markedly in their evolutionary dynamics. While receptor genes are reported to undergo rapid evolution, the evolutionary forces acting on signalling and effector genes remain more debated (Sackton, et al. 2007; Hanson, et al. 2016). Sackton et al (2007) showed that signalling genes with modulatory functions are subject to positive or ‘diversifying’ selection (i.e. selection driving fixed differences among species), whereas effector genes tend to turnover in gene number faster. However, subsequent studies, including those in other organisms, have found that effector genes can evolve rapidly by both gene duplication and positive selection (Tennessen 2005; Hollox and Armour 2008; Hanson, et al. 2016; Hanson and Lemaitre 2020; Hanson, et al. 2023). In fact, orthologs of some effector genes in these pathways can be difficult to identify due to their high rates of sequence divergence (Sackton and Clark 2009; Hanson, et al. 2016).

Such analyses suggest that antiviral defence mechanisms may be especially prone to experiencing strong selection, as seen in vertebrates (e.g. Enard, et al. 2016; Ito, et al. 2020; Scheben, et al. 2023). Notably, RNAi pathway components are among the most rapidly evolving genes in *D. melanogaster* and closely related species, particularly those involved in small RNA biogenesis and antiviral defence (Obbard, et al. 2006; Obbard, Gordon, et al. 2009; Palmer, et al. 2018; but compare Hill, et al. 2019). Similar patterns have recently emerged for the cGAS-STING pathway in *Drosophila*, where species encode variable numbers of cGLRs (cGAMP-like receptors), often exhibiting species-specific expansions and functional divergence (Cai, et al. 2023). For example, different cGLR-generated cyclic dinucleotides (Such as 2′3′-cGAMP, 3′2′-cGAMP, 2′3′-c-di-AMP) vary across species in their ability to inhibit *Drosophila* C virus (DCV), suggesting a lineage-specific functional diversification (Cai, et al. 2023).

However, most insights into the factors shaping the drosophilid immune system have come from a handful of species and/or immune genes, and their generality remains unclear. Specifically, comparisons limited to *D. melanogaster* and its close relatives have likely captured only a subset of immune adaptation, leaving broader patterns unexplored. Comparative analyses across more distantly related clades remain scarce, but one study found rapid evolution of Toll signalling genes in *D. innubila*, contrasting with what was found for *D. melanogaster*, where RNAi pathway genes show the fastest rates of evolution (Hill, et al. 2019). Broad phylogenetic sampling across the Drosophilidae now offers a unique opportunity to quantify how immune gene families evolve over tens of millions of years of divergence, spanning diverse ecological niches, microbial exposures, and life histories (Kim, et al. 2024; Dhakad, et al. 2025). Early analyses of a small number of immune genes already suggested lineage-specific adaptation between the *melanogaster* and *virilis* groups, suggesting that different pathogen pressures may shape distinct evolutionary trajectories (Morales-Hojas, et al. 2009). Expanding both the number of species and the immune gene repertoire should improve power to detect lineage-specific adaptations and can reveal macroevolutionary trends obscured in narrow comparisons (Lažetić and Troemel 2021).

Although numerous studies have identified high rates of nonsynonymous substitution or gene family turnover in immune genes, relatively few have had the power to simultaneously or systematically co-analyse additional molecular or functional features that might explain variation in evolutionary trajectories. Importantly, the evolutionary rate of a proteins may be influenced only weakly by the functional role of the protein, and other factors, such as gene expression level, protein structure, gene length, intron number, recombination rate and protein-protein interaction may dominate, when they are taken in account (Drummond, et al. 2005; Larracuente, et al. 2008; Zhang and Yang 2015; Hagai, et al. 2018; Moutinho, et al. 2019; Zhong, et al. 2021). For example, genes with higher baseline expression levels tend to evolve more slowly, likely due to pleiotropic constraints and costs of misfolded proteins (The expression level-evolutionary rate anticorrelation; Drummond, et al. 2005; Hagai, et al. 2018; Zhong, et al. 2021). Similarly, genes encoding structurally ‘buried’ or interaction-rich proteins often show stronger purifying selection due to higher functional constraint (Moutinho, et al. 2019; Chaurasia and Dutheil 2022). By contrast, proteins with high relative solvent accessibility (RSA)—that is, those with many surface-exposed residues—may offer more opportunities for adaptive substitutions, particularly in immune receptors and effectors interacting directly with pathogens (Moutinho, et al. 2019). A gene’s number of physical or genetic interactions may constrain its evolutionary flexibility, as highly connected genes can have wider systemic effects (Pang, et al. 2010; Papakostas, et al. 2014; Zhang and Yang 2015). Likewise, gene length may affect both mutation target size and potential for functional modularity, which could differentially shape immune versus non-immune gene evolution (Larracuente, et al. 2008; Zhang and Yang 2015; Moutinho, et al. 2019). Integrating these potentially important, but often ignored, predictors into analyses of immune gene evolution could provide a more mechanistic understanding of why some genes or sites evolve adaptively, while others do not.

To address this, we examine the evolutionary dynamics of immune gene families across 304 species of Drosophilidae. Using a curated set of immune-related genes and length- and location-matched non-immune genes (‘controls’), we first estimate relative rates of protein sequence divergence (dN/dS), the proportion of sites under diversifying selection, test for evidence of positive selection, and quantify gene family turnover (λ). Then, using a mixed-model meta-analytic approach, we analyse these estimates to quantify the role of structural and regulatory gene features—such as relative solvent accessibility (RSA), baseline gene expression, number of gene interactions, and gene length—in predicting variation in evolutionary outcomes, regardless of immune function. At the same time, we test whether immune genes differ systematically from non-immune genes in their rates of evolution, while accounting for the gene-level predictors. Finally, we ask whether different immune classes and pathways—such as ‘recognition’, ‘signalling’, and ‘effector’ genes, or Toll, Imd, and RNAi pathways—exhibit distinct patterns of sequence divergence and gene turnover. Together, this study aims to reveal the general determinants of rapid protein evolution and assess how functional roles shape the tempo and mode of immune gene diversification.

## Results and Discussion

To investigate the evolutionary dynamics of immune gene families across Drosophilidae, we analyzed a curated set of *D. melanogaster* immune genes and their homologs in 304 species, alongside size- and location-matched non-immune genes (‘controls’). We estimated rates of protein sequence evolution using codon-based alignments and gene trees to compute gene-wide dN/dS values, and to detect signals of episodic diversifying selection (i.e. positive selection driving divergence among species), using the approach implemented in HyPhy ‘BUSTED’ (Murrell, et al. 2015). We estimated gene family turnover rates (duplication and loss per million years) using CAFE5 (Mendes, et al. 2021), under a stochastic birth-death model calibrated to an approximately-dated relaxed-clock species phylogeny.

To identify the factors shaping immune gene evolution, we fitted a series of Bayesian multivariate generalized linear mixed models using the MCMCglmm R package (Hadfield 2010). These models simultaneously inferred the predictors of five evolutionary responses: the log-transformed dN/dS ratio, log-transformed gene turnover rate (λ), a binary indicator of whether a gene family exhibits nonzero turnover, the proportion of sites (codons) inferred to evolve under diversifying selection, and the presence or absence of statistically ‘significant’ evidence for episodic selection (see Material and Methods). We included four fixed effect predictors based on previously reported determinants of the rate of adaptive evolution, including estimated relative solvent accessibility (RSA), baseline gene expression in *D. melanogaster*, number of gene interactions, and gene length (Drummond, et al. 2005; Larracuente, et al. 2008; Zhang and Yang 2015; Moutinho, et al. 2019; Chaurasia and Dutheil 2022). This integrated framework allowed us to test which gene features best predict patterns of sequence evolution and gene turnover, and whether immune genes exhibit distinct evolutionary dynamics compared to non-immune ‘control’ genes, after accounting for gene-level predictors. We further examined how these patterns differ across immune functional classes and immune pathways, while statistically controlling for gene-level structural and regulatory constraints.

### Structural and gene-level features predict patterns of molecular evolution and turnover

In our analysis, relative solvent accessibility (RSA) emerged as the strongest overall predictor of molecular adaptation. Genes encoding proteins with more surface-exposed residues (i.e., higher RSA) showed markedly elevated rates of sequence evolution: dN/dS increased by 17.2% per standard deviation in RSA (CI [13.0, 21.0], p<0.001), and the proportion of sites under diversifying selection increased by 24.2% (CI [16.5, 32.1], p<0.001). RSA also predicted the probability of detecting episodic selection, with a 3.4% increase in probability per SD (CI [1.7, 5.0], p < 0.001). These findings align with studies that integrated structural and evolutionary analysis, showing that surface-exposed residues evolve faster due to weaker structural constraint and increased functional accessibility (Moutinho, et al. 2019; Chaurasia and Dutheil 2022). In contrast, RSA was not significantly associated with gene turnover rate (λ) among those families that showed some copy number variation (p = 0.35). However, RSA was negatively associated with the probability of any gene family size variation, showing a 3.6% decrease per SD (CI [-5.2, –2.0], p<0.001). This suggests a potential trade-off; while surface-exposed residues are more likely to undergo adaptive sequence change, the genes encoding such proteins are less prone to duplication or loss, possibly due to dosage sensitivity and functional pleiotropy.

Gene length is well known to correlate with rates of evolution, with longer genes evolving more slowly than shorter ones (Yang and Gaut 2011; Zhang and Yang 2015; Soni and Eyre-Walker 2022). Consistent with this, we found an increase in 1kb of gene length is associated with a 4.9% reduction in dN/dS (95% HPD CI [-7.1, -2.7], p<0.001) and a ∼9% decrease in proportion of sites under diversifying selection (CI [-12.7, -5.2], p<0.001; see Supplementary file 1 for model summary). Several mechanisms may underlie this pattern, including more potential sites for deleterious mutations to occur, increased interference from linked selection constraining the efficacy of natural selection (Hill-Robertson effects), greater functional constraint due to more interaction partners or regulatory complexity in longer genes, and the potential for faster protein synthesis in shorter genes (Loewe and Charlesworth 2007; Larracuente, et al. 2008; Zhang and Yang 2015; Moutinho, et al. 2019). Interestingly, despite showing reduced dN/dS and fewer positively selected sites, longer genes were *more* likely to be inferred as evolving under episodic diversifying selection (as assessed by the BUSTED model), with each 1 kb increase in gene length associated with a 16% increase in the probability of detecting such selection (p<0.001). This association may reflect the increased statistical power to detect selection in longer genes, which provide more codon sites where diversifying selection could be detected even if most of the gene evolves under purifying or neutral selection (Yang and Dos Reis 2011; Murrell, et al. 2015). In contrast, gene length was not significantly associated with the rate of gene turnover (λ) among genes showing any variation in family size (p = 0.57; Supplementary file 1). However, longer genes were significantly less likely to exhibit any family size variation at all, with a 1.9% decrease in the probability of observing any turnover, per additional kilobase of coding sequence (p = 0.002). Together, these results suggest that longer genes are not only more constrained at the sequence level, but also less prone to gene duplication and loss, reinforcing the notion that gene length imposes broad constraints on both molecular and genomic evolution.

Gene expression level, widely considered a major constraint on protein evolution (Drummond, et al. 2005), was associated with measures of sequence evolution but not copy number variation. As expected, genes with higher baseline expression in *D. melanogaster* showed a 1.3% decrease in *dN/dS* per expression level (CI [-1.6, -0.1], p<0.001) and a 1.4% reduction in the proportion of sites under diversifying selection per expression level (CI [-2.0, -0.08], p<0.001). These findings are consistent with previous observations in which highly expressed genes experience stronger purifying selection, likely due to constraints imposed by protein misfolding, translational error sensitivity and high pleiotropic effects (Drummond, et al. 2005; Zhang and Yang 2015; Bédard, et al. 2022). However, we also detected a modest positive association between expression level and the probability of detecting episodic selection, with 0.4% increase per expression level (p<0.001). This suggests that even highly expressed genes can be targets of occasional bursts of adaptive evolution, potentially reflecting context-dependent pressures—for example, tissue-specific or inducible expression under stress or infection (Gu and Su 2007; Larracuente, et al. 2008; Kryuchkova-Mostacci and Robinson-Rechavi 2015; Stanley and Kulathinal 2016).

As expected, genes with a higher number of genetic/protein interaction partners were under stronger evolutionary constraint. Specifically, each additional interaction reported in *D. melanogaster* was associated with a 0.41% decrease in dN/dS (CI [-0.51, -0.31], p<0.001) and a 0.44% reduction in the proportion of sites under diversifying selection (CI [-0.62, - 0.26], p<0.001). These findings are consistent with the ‘centrality–lethality’ hypothesis, which posits that highly connected proteins (network hubs) evolve more slowly due to their essential roles and pleiotropic effects on multiple cellular processes [Fraser et al (2002), Hahn et al (2005) but see also Jordan et al (2003) and Mekic et al (2024)]. In contrast, interaction count had no significant effect on gene turnover rates among variable families (p = 0.27), nor on the likelihood of family size variation (p = 0.87).

### Immune genes exhibit elevated sequence divergence, but lower turnover compared to non-immune genes

After accounting for gene length, baseline expression, number of genetic/protein interactions, and relative solvent accessibility (RSA), we found that immune genes do evolve significantly faster than non-immune genes at the protein sequence level, but exhibit lower rates of gene turnover (Fig. 1). Immune genes had a mean dN/dS of 0.10, compared to 0.07 for non-immune genes—both slightly higher than the estimates from *D. melanogaster* and its closest relatives reported by Sackton et al (2007; median dN/dS = 0.08 and 0.06). The 23% higher dN/dS of immune genes (95% HPD CI [13.8, 32.7], p<0.001; Fig. 1A; Supplementary file 1) was reflected in elevated signatures of positive selection. Immune genes had a 26.3% higher proportion of sites reported to evolve under diversifying selection (CI [8.1, 44]; p=0.001; Fig. 1B) and an 8.8% increase in the probability of detecting episodic selection (CI [3.2, 14.6], p<0.001), indicating more rapid adaptive evolution than non-immune genes. These results are consistent with previous findings that immune genes are among the most rapidly evolving in many taxa (Viljakainen, et al. 2009; Harpur and Zayed 2013; Scheben, et al. 2023), including *Drosophila* (Sackton, et al. 2007; Obbard, Welch, et al. 2009; Shultz and Sackton 2019).

**Fig. 1:**
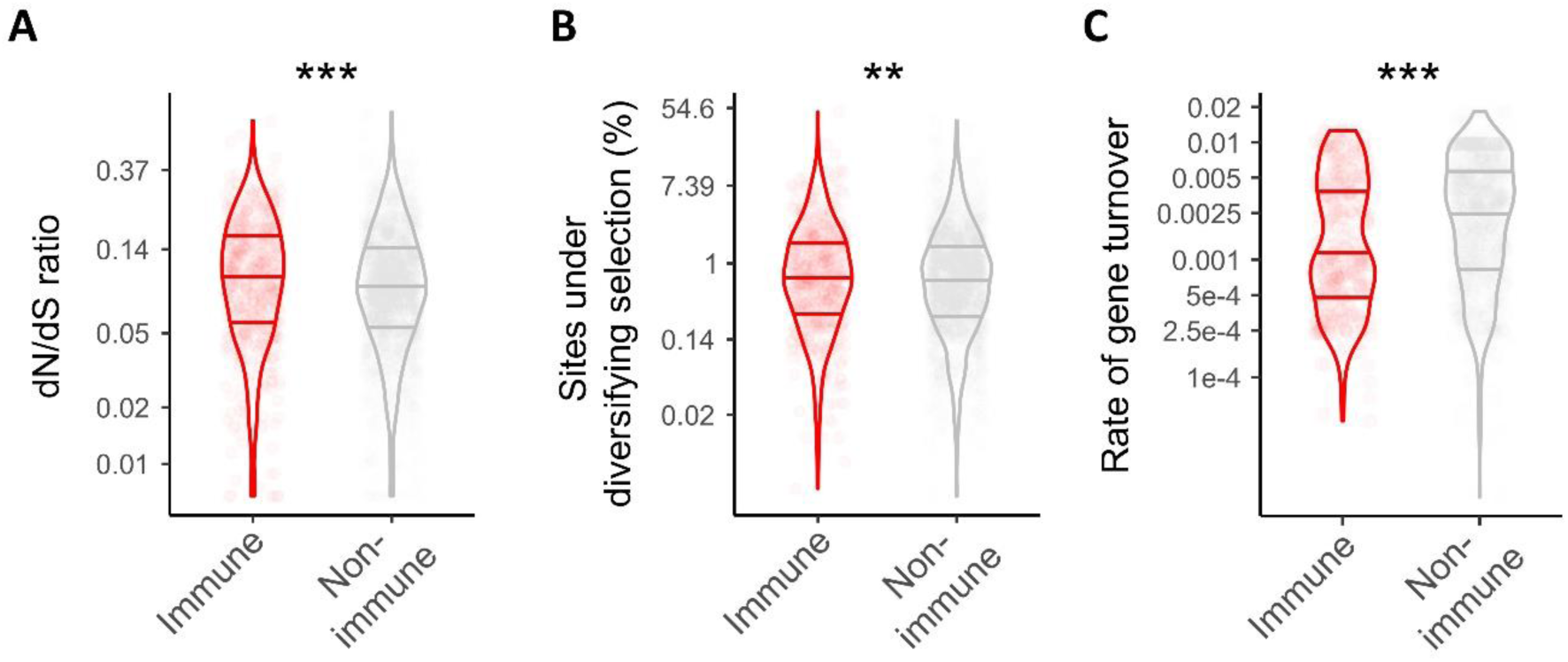
Comparative evolutionary rates of immune and non-immune genes across species of Drosophilidae. Violin plots show (A) the non-synonymous to synonymous substitution rate ratio (*dN/dS*), (B) the proportion of sites (codons) under episodic diversifying selection (as inferred from BUSTED), and (C) the estimated rate of gene turnover (λ, births and deaths per million years). Only, λ values > 3 × 10^-^*^7^* included (others were considered to be zero, and fitted as a binary outcome). The y-axes are plotted on a log scale for visualization but labelled with untransformed values. Immune genes (red) show significantly higher dN/dS and a greater proportion of sites under positive selection, but lower turnover rates, compared to position- and size-matched non-immune controls (grey). The plotted points depict raw estimates for each HOG, not GLMM model fits, but ‘significance’ levels are derived from the Bayesian MCMCglmm models that also include gene length, expression level, number of interactors, and RSA as fixed effects: p < 0.05 (*), p < 0.01 (**), p < 0.001 (***).

Across the Drosophilidae as a whole, immune genes were also more likely to exhibit detectable variation in gene family size, with a 6.2% increase in the probability detecting some variation in copy number (CI [2.4, 10], p = 0.001; Supplementary file 1). However, among those genes that did display variation in gene family size, immune genes had a significantly *lower* estimated turnover rate (λ = 0.003) compared to non-immune genes (λ = 0.005), a 44.2% reduction (CI [-55.2, -32.5]; p<0.001; Fig. 1C; Supplementary file 1). This finding is somewhat unexpected, as many previous studies have emphasized the role of gene duplication and loss in immune gene evolution (Sackton, et al. 2007; Salazar-Jaramillo, et al. 2014b; Levine, et al. 2016; Crysnanto and Obbard 2019). However, these studies were restricted to a small subset of *Drosophila* species, where recent duplications and losses are more visible, whereas over deeper evolutionary timescales immune gene families may be relatively stable, punctuated by lineage-specific bursts of expansion and contraction.

### The role of functional class in immune gene evolution

To examine how evolutionary dynamics vary across immune gene functions, we grouped immune genes into six functional classes: ‘receptors’, ‘effectors’, ‘signalling’, ‘antiviral’, genes involved in ‘multiple’ roles, and other ‘unclassified’ immune-related genes. We found substantial heterogeneity in evolutionary rates among these functional classes. Compared to non-immune genes, several immune classes exhibited elevated dN/dS values. Receptors, effectors, antiviral, and unclassified genes all showed significantly higher dN/dS than non-immune genes (mean dN/dS = 0.07 vs mean dN/dS: receptor = 0.11, effector = 0.12, antiviral = 0.13, all comparisons p<0.001; Fig. 2A; Supplementary file 2). Among immune classes, antiviral proteins evolved most rapidly, with significantly higher dN/dS than signalling proteins (mean dN/dS = 0.13 vs 0.07, p<0.001), consistent with earlier studies highlighting the rapid evolution of antiviral pathways genes (Fig. 2A; Obbard, et al. 2006; Obbard, Welch, et al. 2009; Palmer, et al. 2018). There was also significant variation among immune classes, for example effectors (including AMPs) and receptors also had higher dN/dS than signalling proteins (mean difference from ‘signalling’ for both = 0.04, p < 0.001), in line with their direct roles at the host–pathogen interface resulting in high substitution rates (Fig. 2A; Lazzaro 2005; Sackton, et al. 2007; Unckless and Lazzaro 2016a). In contrast, signalling proteins dN/dS was the lowest among immune classes, and indistinguishable from non-immune proteins (mean dN/dS = 0.073 vs 0.074, p = 0.85), likely reflecting their involvement in conserved pathways with pleiotropic functions (Yu, et al. 2022). The proportion of sites under detectable ‘diversifying’ (i.e. positive) selection mirrored some, but not all, of the trends seen for dN/dS. Effectors and unclassified genes had a significantly higher proportion of sites under diversifying selection compared to non-immune genes (Fig. 2B). Surprisingly, antiviral genes did not differ from non-immune genes in the proportion of positively selected sites, despite their high overall dN/dS (Supplementary file 2). This pattern may reflect weaker purifying constraint on antiviral genes, inflating overall dN/dS without producing strong, recurrent signals of site-specific adaptation.

**Fig. 2:**
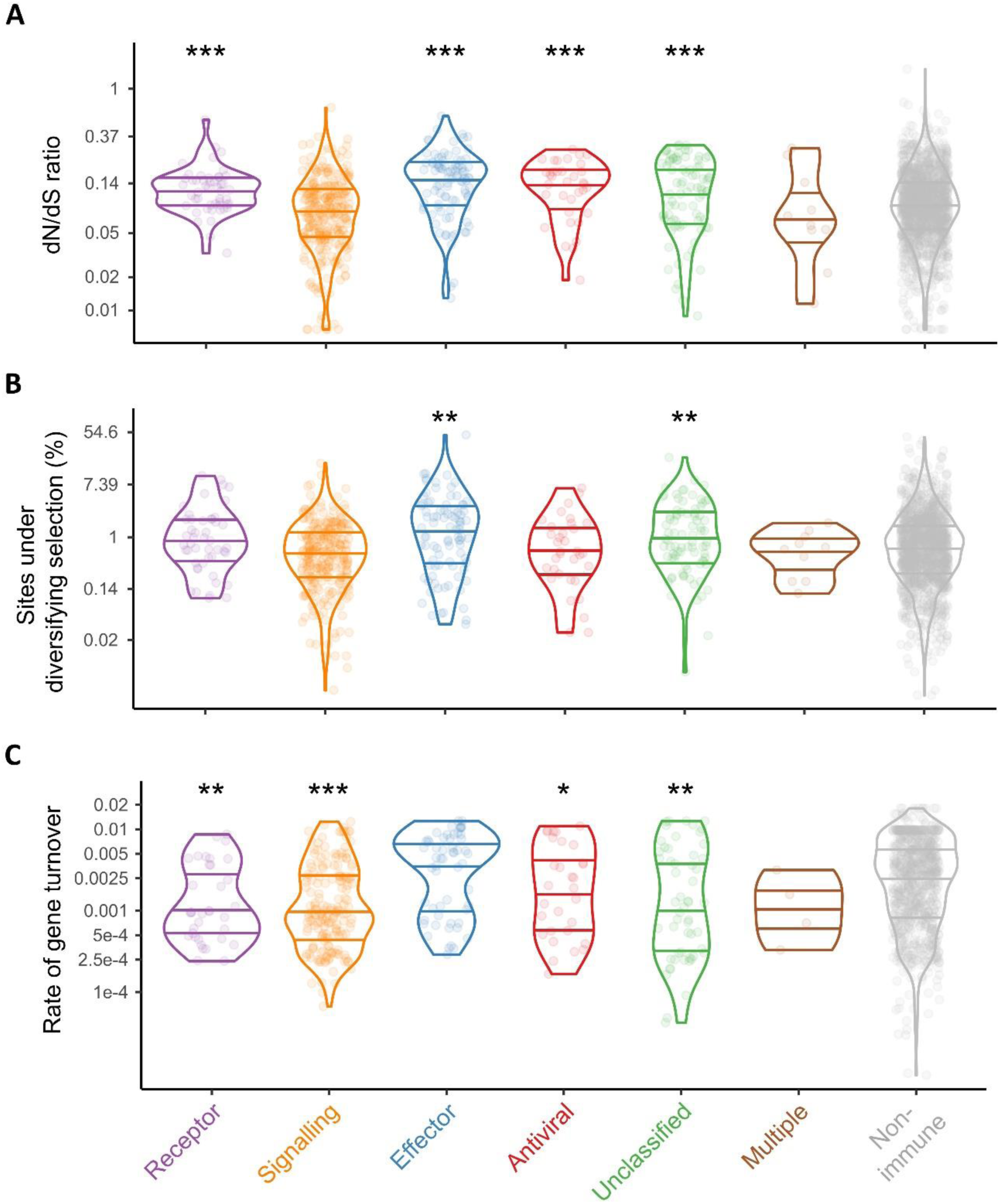
Variation in evolutionary rates among immune functional classes. Violin plots show (A) *dN/dS*, (B) proportion of sites (codons) under episodic diversifying selection, and (C) gene turnover rate (λ, per million years) for receptors (purple), signalling components (orange), effectors (blue), antiviral genes (red), unclassified immune genes (green), and genes assigned to multiple functional categories (brown), compared to non-immune controls (grey). Only, λ values > 3 × 10^-^*^7^* included (others were considered to be zero and fitted as a binary outcome). All y-axes are plotted on a log scale but labelled with untransformed values. The plotted points depict raw estimates for each HOG, not GLMM model fits, but ‘significance’ levels are derived from the Bayesian MCMCglmm models that also include gene length, expression level, number of interactors, and RSA as fixed effects: p < 0.05 (*), p < 0.01 (**), p < 0.001 (***). The p-values are reported with respect to non-immune genes.

Gene turnover dynamics also varied across functional classes, though in less clear-cut ways. Relative to non-immune genes, all immune genes classes except effectors and genes with multiple roles had significantly lower rates of gene turnover (Fig. 2C, Supplementary file 2). Among immune genes, effectors exhibited the highest turnover rates, being duplicated or lost faster than signalling (mean λ = 0.006 vs 0.004, p = 0.001), receptor (mean λ = 0.006 vs 0.004, p = 0.006), or antiviral genes (mean λ = 0.006 vs 0.004, p = 0.026). These results highlight different axes of gene evolution in functional classes of immune genes; effector genes appear to diversify through both duplication and coding sequence change, antiviral and receptor genes have predominantly adapted via coding sequence change only and signalling genes are substantially more conserved across both dimensions.

### The role of pathways in immune gene evolution

To determine whether immune pathways (rather than functional ‘classes’, above) have experienced distinct evolutionary pressures across Drosophilidae, we compared the dN/dS ratio, the proportion of sites under detectable diversifying selection, and gene turnover rates across annotated pathways, using a GLMM approach as above. We found substantial variation in evolutionary rate among immune pathways (Fig. 3, Supplementary file 3). The cGAS-STING pathway genes had the highest dN/dS, more than twice that of non-immune genes (mean dN/dS = 0.20 vs 0.07, p = 0.02) and significantly higher than all other immune pathways except Toll, RNAi, and Imd (mean dN/dS: Toll = 0.12, RNAi = 0.10, Imd = 0.08, all p < 0.05; Fig. 3A; Supplementary file 3). Surprisingly, Toll pathway genes—often reported to be more conserved component of insect immunity (Begun and Whitley 2000; Schlenke and Begun 2003; Sackton, et al. 2007)—had marginally higher dN/dS than the Imd pathway (mean diff. = 0.03, CI [0.007, 0.06], p = 0.04) and slightly higher (but not significantly) dN/dS than RNAi genes (p = 0.65). This contrasts with previous reports from *D. melanogaster-D. simulans* lineages, which emphasized rapid evolution in Imd and RNAi (Sackton, et al. 2007; Obbard, Welch, et al. 2009), but is consistent with broader phylogenetic survey (e.g. quinaria group) showing accelerated Toll evolution in mushroom-breeding species (Hill, et al. 2019). These patterns suggest that the relative rate of pathway evolution may be shaped by lineage-specific pathogen pressures. Pathways involved in broader cellular signalling, such as MAPK and genes classified in “multiple” immune pathways, had the lowest dN/dS (0.06 and 0.07, respectively), consistent with their additional functions in conserved developmental and metabolic processes (Shilo 2014). Despite these differences in overall dN/dS, the proportion of codon sites evolving under detectable episodic diversifying selection did not vary markedly among pathways (Fig. 3B, Supplementary file 3).

**Fig. 3:**
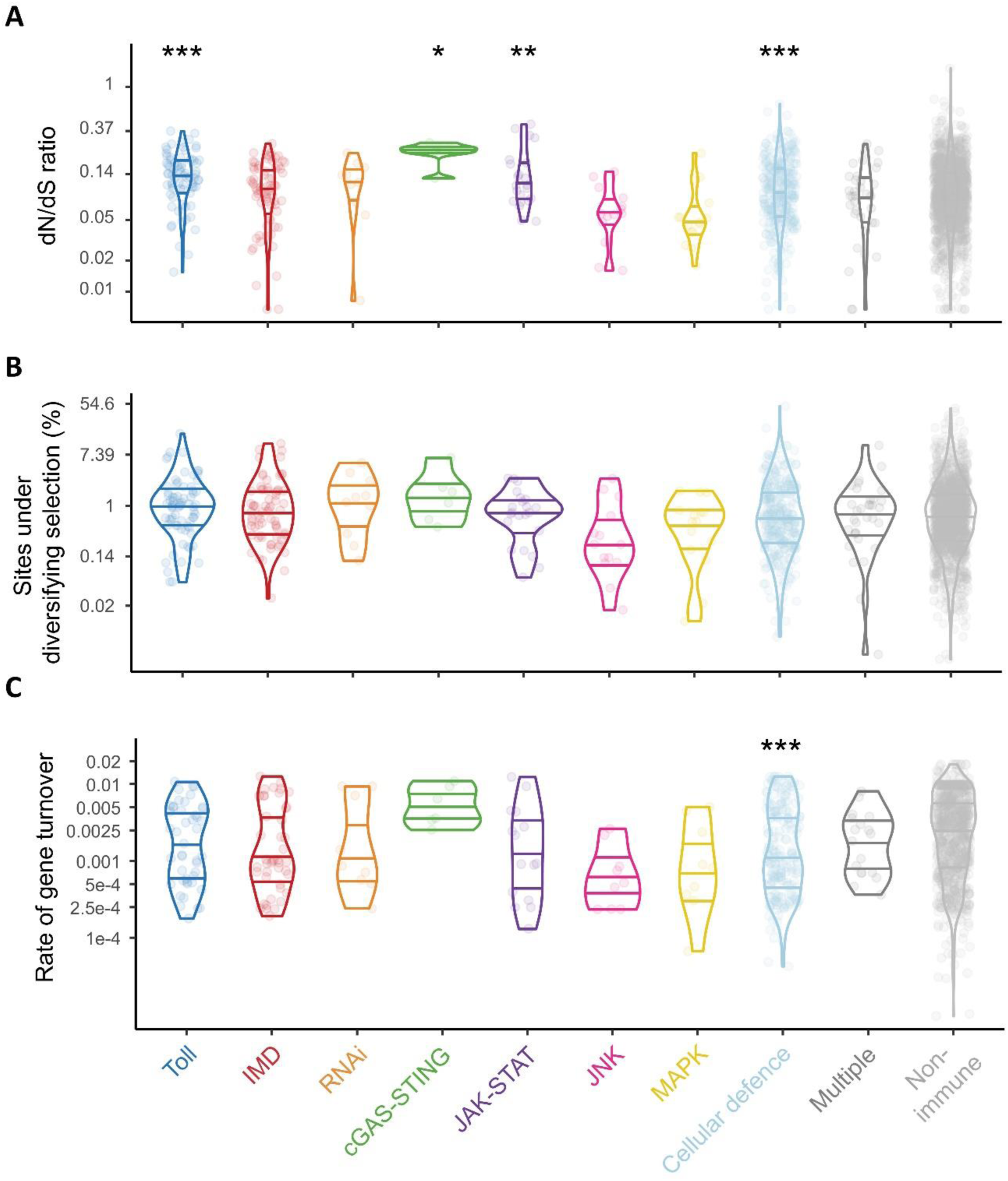
Variation in evolutionary rates among immune pathways. Violin plots show (A) *dN/dS*, (B) proportion of sites (codons) under episodic diversifying selection, and (C) gene turnover rate (λ, per million years) for genes assigned to major immune pathways, compared to position- and size-matched non-immune controls. Only, λ values > 3 × 10^-^*^7^* included (others were considered to be zero and fitted as a binary outcome). All y-axes are plotted on a log scale but labelled with untransformed values. The plotted points depict raw estimates for each HOG, not GLMM model fits, but ‘significance’ levels are derived from the Bayesian MCMCglmm models that also include gene length, expression level, number of interactors, and RSA as fixed effects: p < 0.05 (*), p < 0.01 (**), p < 0.001 (***). The p-values are reported with respect to non-immune genes.

Gene turnover rates were also similar among pathways, except for cellular defence genes, which had significantly lower turnover rate than non-immune genes (mean λ = 0.002 vs 0.005, p < 0.001; Fig. 3C, Supplementary file 3). This stability may reflect functional constraints on phagocytic and encapsulation-related genes, where dosage balance and structural complexity could limit the retention of duplicates. Supporting this idea, a comparative study of haemopoiesis pathway genes and those differentially expressed during the encapsulation response found that haemopoiesis-associated genes are highly conserved and present in *Drosophila* species, regardless of their resistance phenotype (Salazar-Jaramillo, et al. 2014b). Such conservation suggests that the core machinery of cellular immunity evolves under strong stabilizing selection, even across lineages facing diverse pathogen pressures, in contrast to recognition and effector roles, where copy number changes are more frequent.

### Some individual genes may be hotspots of rapid adaptive evolution

Regardless of pathway or role, some immune genes may be targets of recurrent strong selection, i.e. potential ‘coevolutionary hotspots’ (Jiggins and Kim 2007; Obbard, Welch, et al. 2009). Interestingly, dN/dS and gene turnover were uncorrelated after accounting for gene-level predictors (Supplementary figure 1), suggesting that protein sequence divergence and copy number dynamics represent largely independent axes of immune gene evolution. Now, to identify immune genes exhibiting distinct evolutionary trajectories—favouring sequence-level adaptation, gene copy number change, or both—irrespective of their functional class or pathway, we examined genes with exceptionally high dN/dS, gene turnover rate (λ), and the proportion of sites (codons) under episodic diversifying selection (Fig. 4). Genes in the top 2.5% for each metric were considered (i.e., dN/dS ratio >0.25, λ >0.107 gain/loss per million years per gene, >6.7% of sites under episodic diversifying selection). In total, 32 genes were in top 2.5% of at least one metric, of which genes involved in cellular defence were 20, followed by those in the Imd (5), Toll (4), JAK–STAT (2), and cGAS–STING (1) pathways. Only two genes, *Srg1* (Sting-regulated gene 1) and *CG9989* ranked in top 2.5% for both dN/dS and λ (Fig. 4A). Three other genes, *bam* (bag of marbles), *CG14957*, and *CG6357* ranked in top 2.5% for both dN/dS and the proportion of positively selected sites (Fig. 4B). The *CG9733* was the only gene in the top 2.5% for both λ and proportion of selected sites (Fig. 4C). Strikingly, RNAi pathway genes (*Ago2*, *R2D2*, and *Dcr2*) did not rank in the top 2.5% of any metric, despite previous reports of rapid RNAi evolution in *D. melanogaster* and close relatives (Fig. 4; Supplementary file 4; Obbard, et al. 2006; Obbard, Gordon, et al. 2009; Obbard, Welch, et al. 2009; Palmer, et al. 2018). This suggests that such acceleration may be lineage-specific, rather than pervasive across Drosophilidae (Hill, et al. 2019). Likewise, all functional immune classes were equally represented among fastest evolving immune genes, except for ‘antiviral’ class (only one gene). This suggests that while antiviral pathway genes do show elevated rates of evolution on average, they do not individually stand out as the fastest evolving elements of the immune system across the whole of the family Drosophilidae (Hill, et al. 2019).

**Fig. 4:**
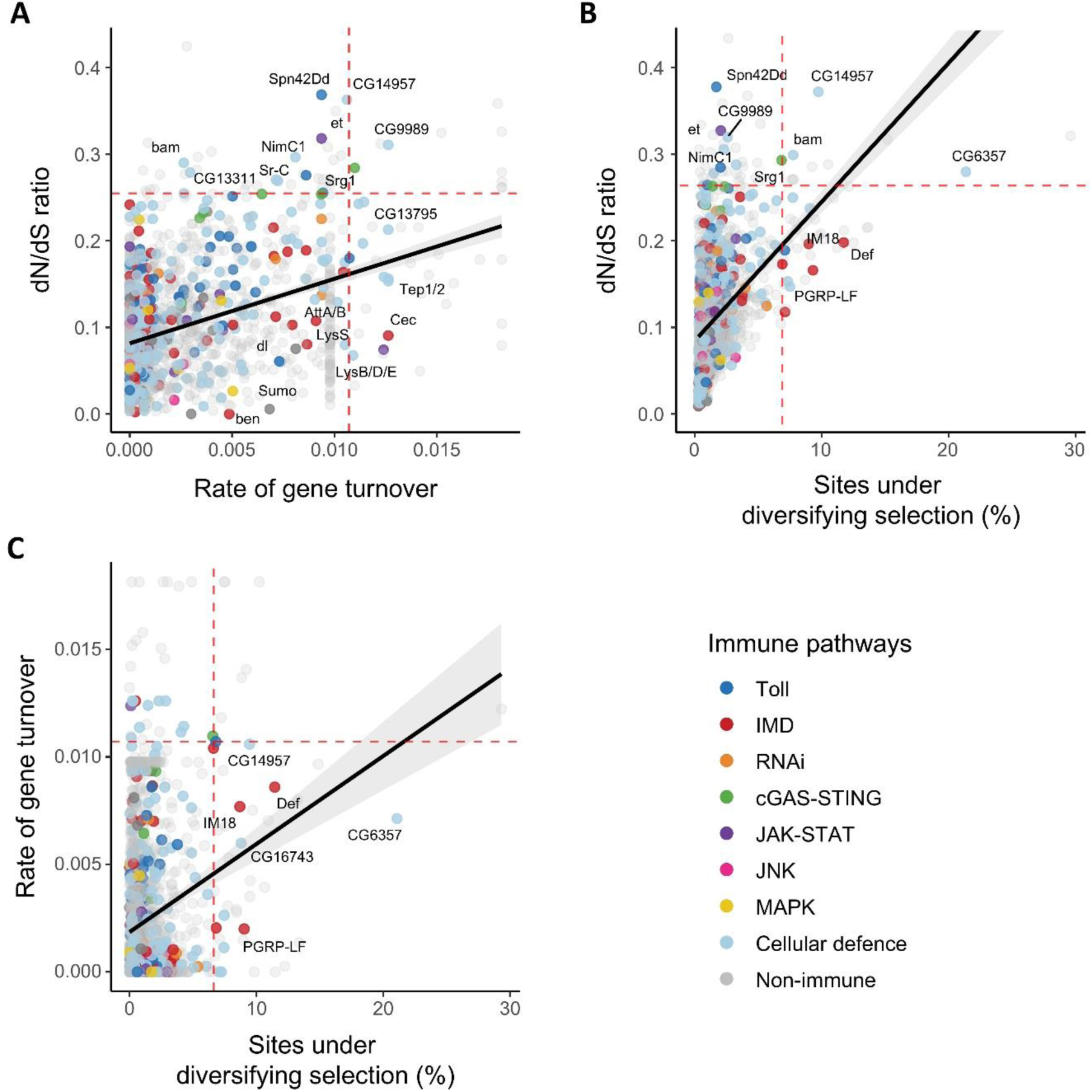
Pairwise relationships among evolutionary metrics for immune genes: (A) dN/dS versus gene turnover rate (λ), (B) dN/dS versus proportion of sites under episodic diversifying selection, and (C) proportion of sites versus gene turnover rate. Red horizontal and vertical lines indicate the top 2.5% thresholds for each metric. Each point represents an immune gene, colored according to its assigned immune pathway.

In contrast to some previous studies in *Drosophila*, we found evidence for adaptive protein evolution in AMPs. For example, *defensin* and *IM18* carried seven and three positively selected sites, respectively, most of which were in the mature functional peptide (5 out of 7 for *defensin* and all sites for IM18, Supplementary figure 2). This supports growing evidence that some AMPs in *Drosophila* are also experiencing strong selection [Unckless et al (2016b), also in other insects Viljakainen et al (2008), Erler et al (2014), Harpur et al (2013)]. Other AMPs, such as *cecropins* and *lysozymes*, instead showed high turnover rates. The *cecropins* in particular have undergone multiple independent expansions and losses across *Drosophila* (Ramos-Onsins and Aguadé 1998; Sackton, et al. 2007). Interestingly, this AMP family is ancient in Diptera but appears to have been lost entirely from the subfamily Steganinae, with only truncated copies recovered in two *Amiota* species (Supplementary figure 3; Hultmark 1993).

Several signalling genes also showed evidence for rapid evolution. *Bam*, *serpins*, *Pten*, *Skanda* and *eyetransformer* (*et*) had high dN/dS, with *bam* also ranking in the top 2.5% for the proportion of positively selected sites (Fig. 4; Supplementary file 4). The gene *bam* is essential for germline stem cell differentiation and gametogenesis and has previously been shown to evolve adaptively in *D. melanogaster* group species, likely driven by *Wolbachia* infections (Flores, et al. 2015; Bubnell, et al. 2022). Our analysis revealed 13 positively selected codons across at least 65 branches of the *bam* gene tree, confirming repeated protein diversification across Drosophilidae (Supplementary figure 2; Bubnell, et al. 2022).

Receptors were also well-represented among the most rapidly evolving genes, including *PGRP-LF* and *PGRP-SB2*, as well as four phagocytic receptors: *Tep*, *Sr-C*, *NimC1*, and *NimB3*. The high representation of phagocytic receptors may suggest that their microbial targets are more labile than the ligands of *PGRPs* or *GNBPs*, driving repeated host–pathogen coevolution. Among these, two families stood out for their particularly strong adaptive signatures: thioester-containing proteins (Tep) and scavenger receptors of class C (Sr-C). Tep genes (*Tep1* and *Tep2*) harboured 148 codon sites under episodic diversifying selection, while Sr-C genes (*Sr-CI*, *Sr-CII*, *Sr-CIII* and *Sr-CIV*) had 111 such sites (Fig. 5A).

**Fig. 5:**
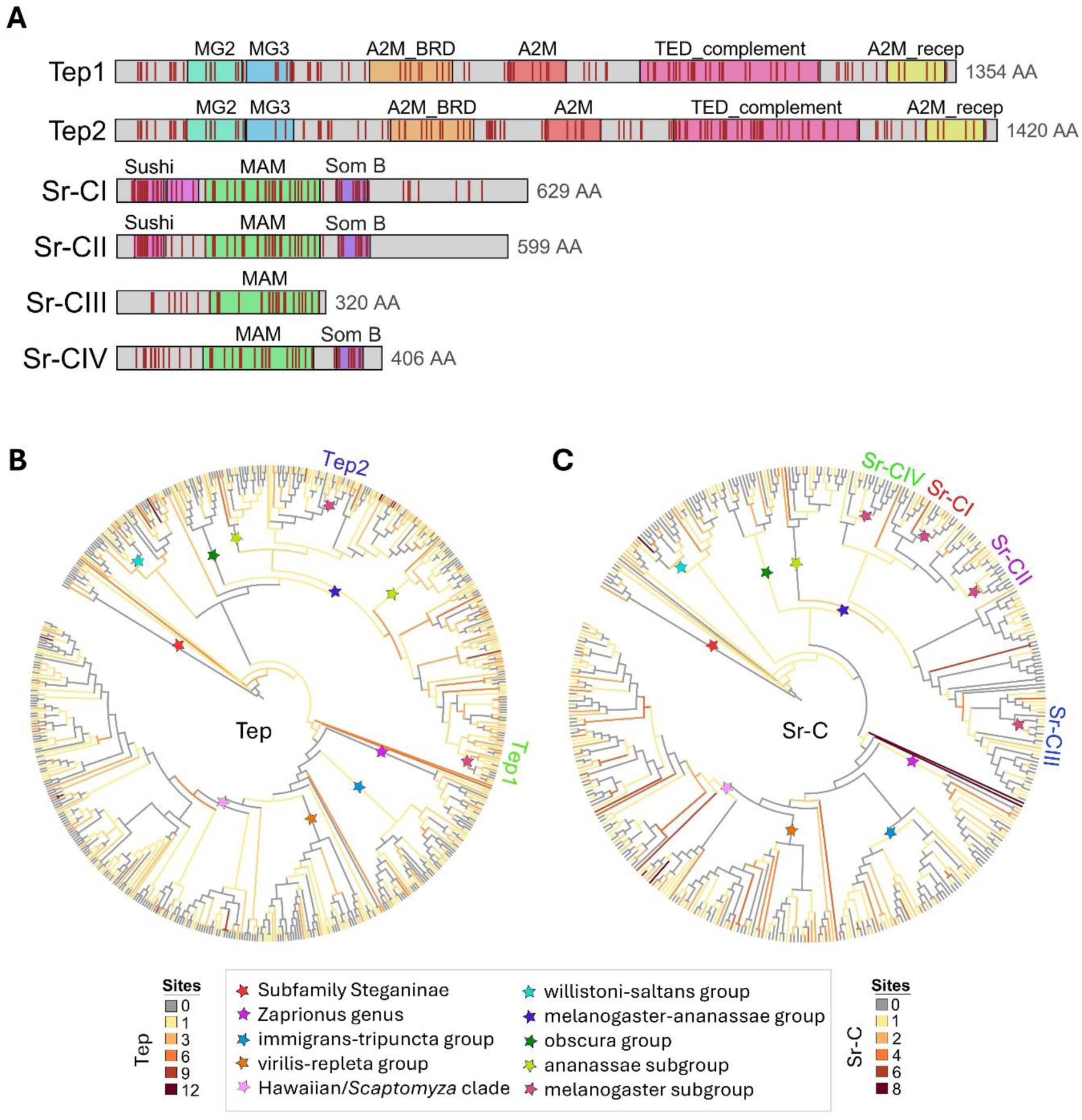
Evolutionary patterns of the thioester-containing protein (Tep) and class C scavenger receptor (Sr-C) families in Drosophilidae. (A) Schematic gene structures of Tep and Sr-C receptors showing conserved functional domains (not to scale) and approximate positions of positively selected sites (red bars). Positively selected sites were identified with MEME. (B, C) Maximum-likelihood gene trees (topology only) of Tep (B) and Sr-C (C) families, rooted by the subfamily Steganinae. Branch colors indicate the number of sites (codons) under episodic diversifying selection (as estimated by MEME). Major *Drosophila* group radiations are marked with stars; *D. melanogaster* paralogs are labelled at the tips.

In Tep proteins, selection sites clustered around conserved MG2/3 (macroglobulin), TED (thiol ester-containing domain), and A2M domains (α-2-macroglobulin; Fig. 5A; Sackton, et al. 2007), which mediate pathogen opsonization and binding to pathogen surface, facilitating their phagocytosis (Williams and Baxter 2014; Shokal and Eleftherianos 2017). In Sr-C proteins, selection sites clustered in Sushi (or Complement control protein, CCP), MAM (meprin/A5-protein/PTPµ) and SomB (Somatomedin B) domains (Fig. 5A; Lazzaro 2005), which are critical for microbial recognition and endocytosis in plasmatocytes (Zani, et al. 2015). Notably, *Tep* genes had markedly higher gene turnover rates (λ = 0.012) than most immune families, reflecting repeated duplication and loss events across the phylogeny. For example, the *D. melanogaster* paralogs *Tep1* and *Tep2* originated before the *melanogaster*-*ananassae* split, whereas other lineages—such as the obscura, willistoni–saltans, repleta–virilis, and Hawaiian and Scaptomyza clades—either retained a single ancestral copy or experienced independent expansions (Fig. 5B). By contrast, *Sr-C* receptors showed relatively stable copy numbers (λ = 0.005) but higher average dN/dS (0.27), consistent with strong, ongoing amino acid divergence in pathogen-binding domains (Fig. 5C). These patterns likely reflect differences in the functional and structural constraints of the two receptor types. Tep’s are soluble opsonins in the haemolymph, where variation in pathogen communities could perhaps favour the gain or loss of paralogs, expanding the repertoire of pathogen-binding specificities. Whereas Sr-C proteins (Sr-CI and Sr-CII) are membrane-bound phagocytic receptors whose turnover may be more constrained by the need to maintain conserved transmembrane and cytosolic domains, that are essential for signalling and endocytosis (Sojo, et al. 2016). As a result, adaptive change in Sr-Cs is more likely to occur through amino acid substitutions at extracellular binding domains rather than through changes in copy number. Together, these gene-level case studies illustrate how immune gene families can adapt through partially overlapping, but distinct, combinations of sequence level adaptation and gene turnover even within a shared functional context.

## Conclusions

Our comparative analysis of 304 Drosophilidae genomes reveals that different axes of immune gene evolution—protein sequence divergence and gene turnover—are shaped by overlapping, but partly independent, sets of predictors. Relative solvent accessibility (RSA) emerged as the strongest positive predictor of rapid or adaptive protein evolution, consistent with the idea that surface-exposed residues, particularly at functional interfaces, are hotspots for adaptive change. In contrast, protein divergence was negatively associated with gene expression level, gene length, and the number of genetic/protein interactions, patterns that match observations across plants, vertebrates, and insects (Duret and Mouchiroud 2000; Zhang and Yang 2015; Moutinho, et al. 2019). These relationships point to constraints imposed by structural and regulatory complexity—indicating that immune gene evolution is shaped not only by host–pathogen arms races but also by general molecular and genomic features. The rate of gene turnover, however, largely unaffected by these gene- and protein-level predictors.

After accounting for gene length, expression level, number of genetic/protein interactions, and RSA, we found that immune proteins evolved faster at the sequence level than non-immune genes, but exhibited lower overall rates of gene turnover. Nevertheless, while these patterns appear robust, several methodological and biological factors need to be considered in their interpretation. First, our turnover estimates from CAFE5 were restricted to gene families present at the origin of Drosophilidae, which may have excluded some of the most recent and volatile multi-copy families. This conservative approach likely underestimates turnover for certain effector gene families, especially those with lineage-specific origins or extreme expansions. Second, our immune gene set contains a relatively high proportion of conserved signalling genes (∼51%) compared to rapidly evolving effectors (∼16%) and receptors (∼8.8%); this composition could reduce the average differences between immune and non-immune categories for both dN/dS and λ. Nevertheless, by jointly considering genomic predictors, structural features, and functional roles, our analysis provides a robust comparative framework for understanding immune system evolution in Drosophila and offers insights that can inform studies of immunity in other taxa.

## Materials and Methods

### Gene selection and orthology assignment

To investigate the evolutionary dynamics of immune gene families, we curated a list of well characterized immune-related genes in *Drosophila melanogaster* from literature searches (De Gregorio, et al. 2001; De Gregorio 2002; Lindmo, et al. 2008; Early, et al. 2017; Troha, et al. 2018; Cai, et al. 2020), including the recent “The *Drosophila* immunity handbook” (Westlake, et al. 2024). Where possible, we assigned each gene to a known functional class (‘recognition’, ‘signalling’, ‘effector’, ‘antiviral’) and immune pathway (‘Toll’, ‘Imd’, ‘RNAi’, ‘cGAS-STING’, ‘JAK-STAT’, ‘MAPK’). Where membership was unclear, or not unique, we assigned immune genes to a ‘Multiple’ or ‘Unclassified’ category. For each immune gene, we then identified up to four non-immune (‘control’) genes that were approximately matched for size and genome location, with the ‘control’ genes required to be protein-coding, located within ±50 kb of the immune gene in the *D. melanogaster* genome, and between 0.5-2 times its length. This approach should help mitigate the impact of local genomic features (e.g. recombination rate, chromatin accessibility—at least to the extent that synteny is maintained with *D. melanogaster* (Felsenstein 1974; Comeron, et al. 2008; Charlesworth, et al. 2009; Cherry 2010; Soni and Eyre-Walker 2022), and reduced the overall computational burden by limiting the number of genes analysed.

Using the *D. melanogaster* references, immune and ‘control’ gene orthogroups were then identified from our recent comparative annotation of 304 Drosophilidae species (Dhakad, et al. 2025). These Hierarchical Orthologous Groups (HOGs) allowed us to compare gene family evolution across deeply diverged lineages in the family Drosophilidae. Some previous studies have chosen to limit their analyses to a subset of closely-related taxa, thereby avoiding a high proportion of noisy, potentially saturated, long branches (e.g. avoiding branches longer than ∼25 million years; Sackton, et al. 2007 used just 6 species in the melanogaster group). However, as almost no lineages in the present dataset lack sampled close relatives (i.e. only 9 of 606 branches longer than ∼25 million years, mean branch length 3.7 million years), such saturation is unlikely to prove problematic.

### Estimating rates of protein sequence evolution

To quantify protein-coding sequence evolution, we used the ‘BUSTED’ model from the HyPhy package (Murrell, et al. 2015). Conditional on gene tree topology, BUSTED tests for ‘episodic’ diversifying selection at any site on any branch of a phylogeny (i.e. selection in favour of amino-acid change on some, but not all, branches). BUSTED returns an overall estimate of the ratio of non-synonymous to synonymous substitutions (dN/dS) per ‘gene’ (Hierarchical Orthogroup; HOG) and reports whether there is evidence for a response to positive selection anywhere in the gene tree (Murrell, et al. 2015). Codon alignments for each gene family (HOG) were generated using MACSE (Ranwez, et al. 2018), a codon-aware aligner specifically designed to handle the frameshifts and sequencing errors common in large, diverse genome-scale datasets. To mitigate the risk of misalignment and subsequent detection of false positives in selection analyses, we applied a series of quality controls in the MACSE OMM pipeline, which uses MAFFT as an aligner to handle larger datasets (Katoh and Standley 2013). First, a pre-filter step to remove long, non-homologous regions that may result from mis-annotation (such as retained introns or gene fusions). Second, HmmCleaner to identify and mask residues that appear misaligned at the amino acid level (Di Franco, et al. 2019). These masked positions were then mapped back to the nucleotide alignment. A third post-processing filtering step was applied to eliminate patchy and isolated codons (sequences with >80% of codons masked were excluded entirely). Finally, we trimmed the 5’ and 3’ extremities of the alignments, discarding poorly aligned ends until a site with at least 70% of nucleotides was reached. These steps collectively aimed to reduce alignment artifacts that could otherwise inflate dN/dS estimates or lead to spurious detection of selection. However, we note that this may come at the cost of excluding rapidly diverging regions and thus reducing the power to detect strong selection. Gene trees were reconstructed for each HOG using IQTREE2 (Minh, et al. 2020), with the best-fit substitution models identified by ModelFinder (Kalyaanamoorthy, et al. 2017) under the Bayesian Information Criterion. We assessed branch support using 1,000 ultrafast bootstrap replicates. These phylogenies and alignments were then supplied to BUSTED for likelihood-based detection of positive selection. BUSTED was run in MPI-parallelized mode with default settings, specifying the entire tree as foreground (to test the entire phylogeny for positive selection) for both immune and non-immune genes. We extracted the gene-wide mean dN/dS values, the proportion of sites detected to evolve under diversifying selection, and a binary indicator as to whether a gene showed evidence of episodic diversifying selection (p-value threshold ≤0.001). For such ‘selected’ HOGs, specific sites undergoing episodic diversifying selection were subsequently identified by HyPhy models ‘MEME’ and ‘FEL’ (Kosakovsky Pond and Frost 2005; Murrell, et al. 2012).

### Estimating gene turnover rate

To estimate the rate of gene turnover (gain and loss per million years, ‘λ’) for immune and non-immune gene families, we used CAFE5 (Computational Analysis of Gene Family Evolution; Mendes, et al. 2021). CAFE models gene family evolution under stochastic birth-death process. However, observed gene copy numbers can be affected by non-biological factors such as assembly errors, variation in genome completeness, or gene family annotation artefacts. To help mitigate this, we first ran the CAFE base model to estimate an error distribution. These error estimates were then incorporated into the subsequent CAFE run to help minimise technical noise prior to estimating ancestral family sizes and λ values (Mendes, et al. 2021). CAFE requires gene families to be present at the root of the species tree. As such, families absent at the root (“orphan” or lineage-specific families) are forced to have an ancestral copy number of at least one. This constraint leads to spurious inferences of gene loss along branches where the family is truly absent, thereby distorting turnover estimates. In addition, we limited the dataset to families with a maximum observed copy number change of 10 genes across all species, as families with extreme variation in size tend to violate model assumptions and often yield unstable or non-converging likelihoods. These filtering steps reduced the dataset to 1,761 ‘gene families’ (i.e. HOGs; 489 immune and 1,272 non-immune HOGs). While these filters were necessary for model accuracy and convergence, they likely biased the analysis toward more conserved families. Specifically, the exclusion of non-rooted families disproportionately removes fast-evolving gene families that arose de novo more recently, or underwent lineage-specific expansions (or families that are lost from an entire ancient clade). As a result, our λ estimates reflect evolutionary patterns among a relatively stable subset of gene families and likely underestimate turnover rates in more dynamic gene categories. Finally, CAFE was run under a gamma model (k=2) with a Poisson distribution for gene family counts at the root to estimate the λ per family. Because CAFE5 does not explicitly return λ = 0, we also recorded a binary variable indicating whether λ was substantially greater than zero (values ≤ 3 × 10^-^*^7^* were treated as effectively invariant). This indicator was included in subsequent linear models to distinguish stable gene families from those exhibiting measurable turnover.

### Mixed-model analyses of sequence evolution and gene turnover

To evaluate how structure, expression level, gene length, number of genetic/protein interactions, and functional properties of genes predict evolutionary dynamics, we took a multivariate ‘meta-analytic’ approach using the Bayesian mixed-model R package MCMCglmm (Hadfield 2010). Our goal was to jointly estimate how gene-level traits such as expression level, relative solvent accessibility (RSA), number of genetic/protein interactions, gene length, and gene function predict the aspects of molecular evolution inferred using HyPhy and CAFE (above).

We fitted a total of 7 different multivariate models (provided in Table 1) that included five response variables for each HOG: (i) the log-transformed dN/dS, (ii) the log-transformed proportion of sites under diversifying selection, (iii) a binary indicator of whether a gene showed evidence of episodic diversifying selection, (iv) the log-transformed rate of gene family turnover (λ), and (v) a binary indicator of whether the estimated turnover rate was clearly distinguishable from zero. These response variables reflect distinct but potentially interrelated evolutionary processes—namely, sequence-level adaptation, copy number evolution, and episodic selection pressure. Considering these traits jointly allowed us to quantify covariation between them and assess whether shared gene properties predict their evolution in parallel. We included fixed effects representing gene-level properties expected to influence evolutionary dynamics: gene length, baseline expression level in *D. melanogaster*, predicted RSA, and number of reported protein and genetic interactions in *D. melanogaster*. Predictors were included in the model as trait-specific fixed effects using the ‘*trait:*’ syntax in MCMCglmm, allowing each explanatory variable to have a separate effect on each evolutionary response (model syntax in Supplementary file 5).

**Table 1:** Summary of multivariate Bayesian models used to investigate evolutionary dynamics of immune gene families. All models included fixed effects for gene length, expression level (FPKM), relative solvent accessibility (RSA), and interaction count; a random effect for positional ID (immune HOG) linking each immune gene with its matched non-immune control (unstructured covariance); and unstructured residual covariance.

| Category | Model | Response Variables | Focal Predictor | Family (link) |
| --- | --- | --- | --- | --- |
| Sequence evolution | M1 | log(dN/dS), log(prop)*, episodic selection (binary) | Gene type (immune vs non-immune) | Gaussian, Gaussian, Threshold (probit) |
|  | M2 | log(dN/dS), log(prop)*, episodic selection(binary) | Immune Class | Gaussian, Gaussian, Threshold (probit) |
|  | M3 | log(dN/dS), log(prop)*, episodic selection (binary) | Immune Pathway | Gaussian, Gaussian, Threshold (probit) |
| Gene turnover | M4 | log($\lambda$), non-zero $\lambda$ (binary) | Gene Type | Gaussian, Threshold (probit) |
| | M5 | log($\lambda$), non-zero $\lambda$ (binary) | Immune Class | Gaussian, Threshold (probit) |
| | M6 | log($\lambda$), non-zero $\lambda$ (binary) | Immune Pathway | Gaussian, Threshold (probit) |
| For Correlation | M7 | log(dN/dS), log(prop)* positive sites, episodic selection (binary), log($\lambda$), non-zero $\lambda$ (binary) | Gene type (immune vs non-immune) | Gaussian, Gaussian, Threshold (probit), Gaussian, Threshold (probit) |

\*Proportion of sites under diversifying selection (on log-scale)

RSA was estimated using predicted structures of *D. melanogaster* proteins obtained from AlphaFold, SWISS-MODEL (Expasy), and ModPipe (Pieper, et al. 2014; Waterhouse, et al. 2018; Abramson, et al. 2024). Solvent accessible surface area (SASA) was computed from the predicted PDB structures using DSSP (https://github.com/PDB-REDO/dssp.git; Kabsch and Sander 1983), and normalized by dividing each residue’s SASA by its reference maximum accessible surface area for a fully exposed residue of that amino acid type (Tien, et al. 2013). Our script for RSA estimation was adapted from Sydykova et al (2018). For each HOG, we calculated the mean RSA across all residues in the corresponding *D. melanogaster* protein. Gene expression was summarized as the mean FPKM (Fragments Per Kilobase of exon per Million mapped fragments) value of each *D. melanogaster* gene across developmental stages and tissues, using expression data obtained from FlyBase (https://flybase.org/). We ranked the HOGs according to mean FPKM values, resulting in 40 expression categories. To quantify protein-protein and genetic interactions, we obtained both protein-protein and genetic interactions for *D. melanogaster* genes from the Drosophila Interactions Database (DroID v2018; Murali, et al. 2011). The median number of interactions across all *D. melanogaster* genes in a given HOG was used as the interaction count for that HOG.

To reduce the impact of any variation associated with genomic location, immune-related genes (i.e. HOGs) were assigned to ‘gene groups’ along with their neighbouring ‘control’ genes, and gene-group ID was fitted as a random effect with an unstructured variance-covariance matrix across traits (analogous to a ‘paired’ test with paired immune and matched non-immune genes). Residual variances were also modelled with an unstructured covariance matrix to capture correlations among traits not explained by the predictors. Our Bayesian priors for fixed effects were chosen to be weakly informative, and a weakly informative parameter-expanded prior was used for the random effects (Hadfield 2010, Supplementary file 5).

To investigate whether immune genes differed from non-immune genes after accounting for structural and regulatory gene-level features, we fitted models (M1 and M4; Table 1) that included gene type (immune or non-immune) as a fixed effect. And, to assess variation within immune genes, we fitted additional models (M2, M3, M5 and M6; Table 1) that included either immune functional class (effector, signalling, recognition, antiviral) or pathway (Toll, Imd, JAK/STAT, RNAi, cGAS-STING) as categorical predictors. All of these models (M1-M7) retained the same structural and regulatory covariates as fixed effects and were fitted separately for either the three sequence evolution statistics (log dN/dS, log proportion of sites under selection, binary episodic selection; models M1-M3) and for the two gene turnover statistics (log λ and binary non-zero λ; Table 1; models M4-M6). In all models, the fixed effects were allowed to vary across traits, enabling us to quantify how immune category membership influences distinct evolutionary processes while adjusting for gene length, expression, RSA, and interactions. All models were run for 2.1 million MCMC steps, with a 100 thousand step burn-in and thinning interval of 100, resulting in a posterior sample of 20 thousand steps. Posterior convergence was assessed visually and through effective sample sizes. Summaries of all models’ variables and predictors are presented in Table 1, and output from each model can be found in Supplementary file 6.

Pairwise contrasts among immune classes or pathways were conducted by subtracting the posterior sample for one level from that of the other, calculating a credible interval for the difference, and estimating ‘pMCMC’ values from the resulting contrasts (i.e., the fraction of the posterior density in the smaller tail overlapping zero). All statistical analysis analyses were performed using R Statistical Software (v4.5; R Core Team 2025) and figures were generated using ggplot2 (Wickham 2016).

## Data availability

All code used for MCMC model fitting, statistical analyses, and figure generation are available at https://github.com/DhakadPankaj/Gene_Family_Evolution. Sequence alignments and gene trees used in this study can be accessed at https://doi.org/10.5281/zenodo.17020656.

## Acknowledgements

We wish to thank Bernard Kim and Dmitri Petrov for making *Drosophila* genome data available to us in advance of their publication, and Jarrod Hadfield for help and advice on the use of MCMCglmm.

