## Supplementary figures and images for "Predictors of protein evolution in the drosophilid immune system"

### Supplementary figure 1

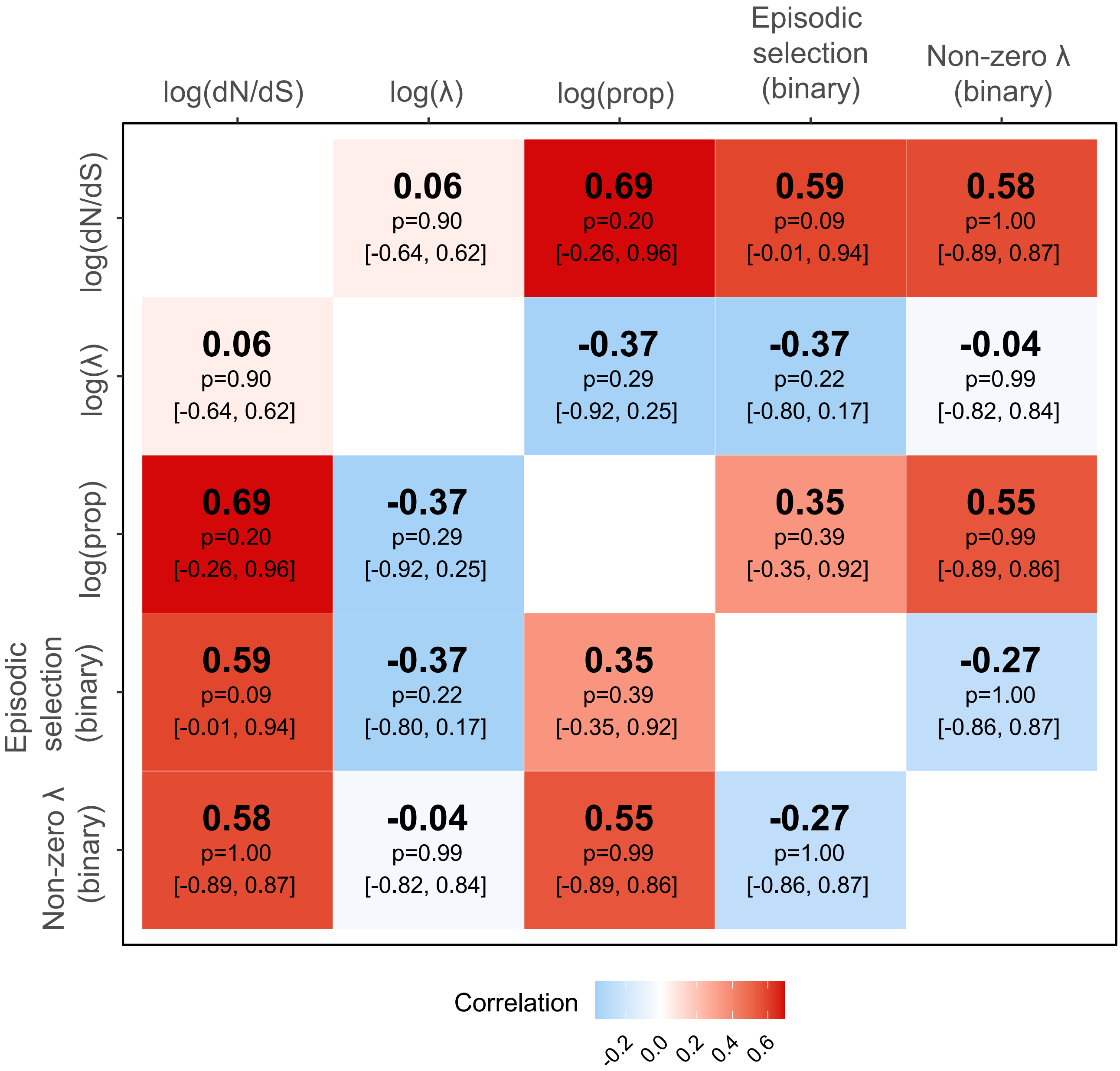

### Supplementary figure 2

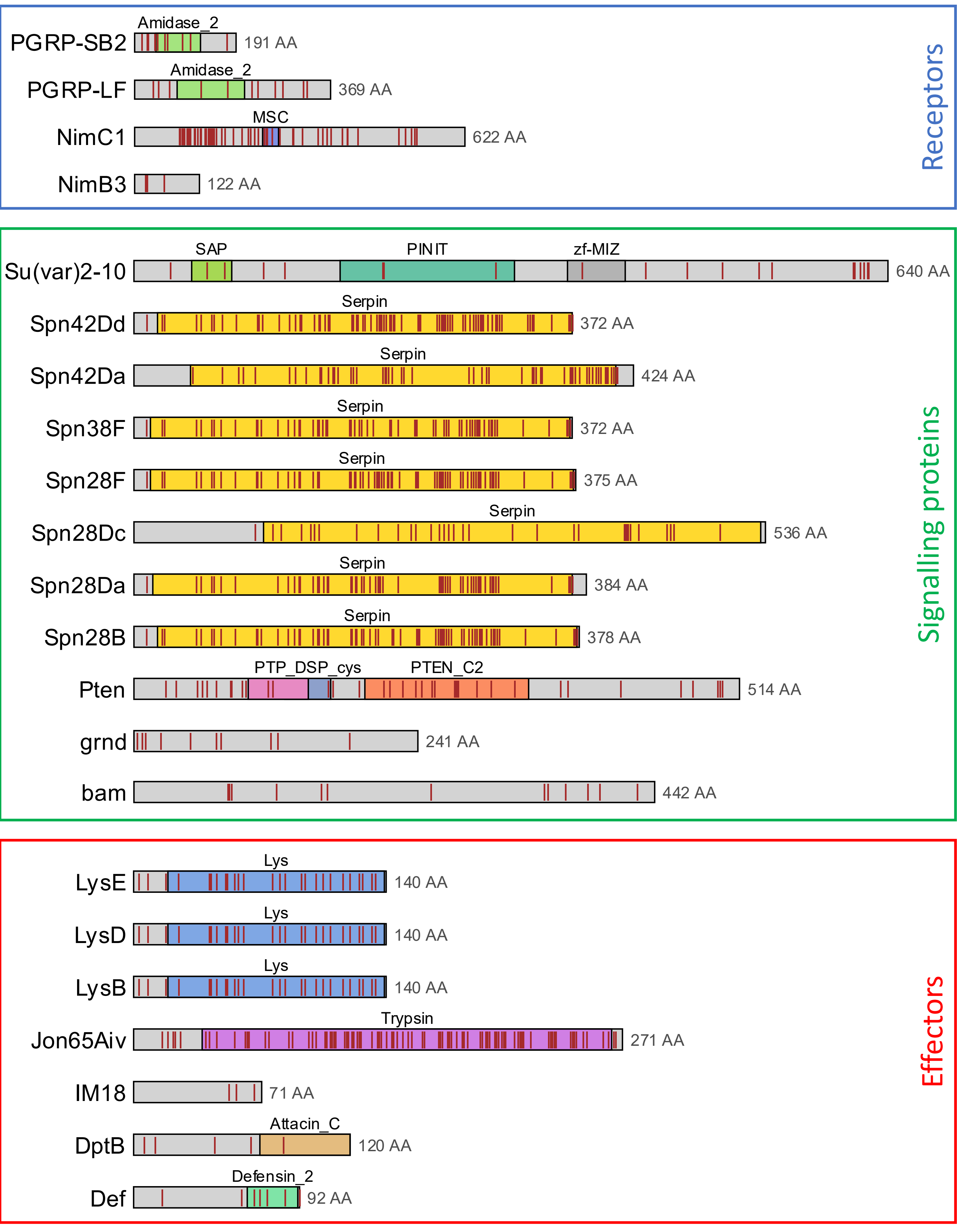

### Supplementary figure 3

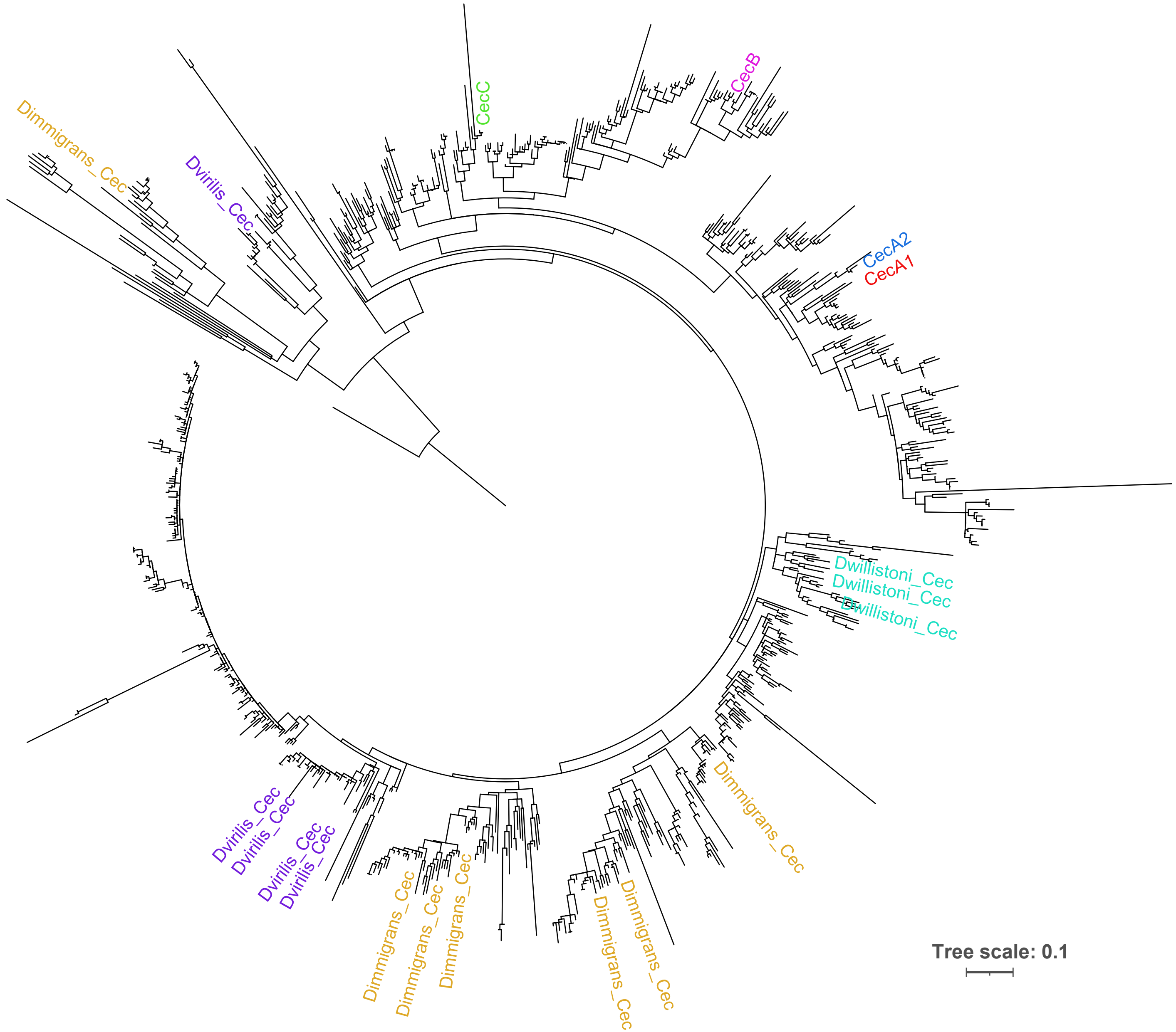
