## Supplementary file 5 for "Predictors of protein evolution in the drosophilid immune system"

### Model syntax and priors for model M1-M7

## 1. M1-M3

```
prior1 <- list(
        B = list(mu = rep(0, 18), V = diag(18) * 1e+10), # Prior for fixed effects
        G = list(G1 = list(V = diag(3), nu = 3, alpha.mu=rep(0,3), alpha.V=diag(3)*1000)),
        R = list(V = diag(3), fix=3, nu = 2.002) # Prior for residual
)

prior2 <- list(
        B = list(mu = rep(0, 33), V = diag(33) * 1e+10), # Prior for fixed effects
        G = list(G1 = list(V = diag(3), nu = 3, alpha.mu=rep(0,3), alpha.V=diag(3)*1000)),
        R = list(V = diag(3), fix=3, nu = 2.002) # Prior for residual
)

prior3 <- list(
        B = list(mu = rep(0, 42), V = diag(42) * 1e+10), # Prior for fixed effects
        G = list(G1 = list(V = diag(3), nu = 3, alpha.mu=rep(0,3), alpha.V=diag(3)*1000)),
        R = list(V = diag(3), fix=3, nu = 2.002) # Prior for residual
)

formula1 <- cbind(log.dNdS, log.prop, busted_positive) ~ trait - 1 + trait:Type +
trait:median_length + trait:FPKM_category + trait:RSA + trait:inter

formula2 <- cbind(log.dNdS, log.prop, busted_positive) ~ trait - 1 + trait:Class +
trait:median_length + trait:FPKM_category + trait:RSA + trait:inter

formula3 <- cbind(log.dNdS, log.prop, busted_positive) ~ trait - 1 + trait:Class +
trait:median_length + trait:FPKM_category + trait:RSA + trait:inter

#Example run
model <- MCMCglmm(
        formula,
        random = ~us(trait):group,
        rcov = ~us(trait):units,
        family = c("gaussian", "gaussian", "threshold"),
        prior = prior,
```

```
        data = df,
        nitt = 2100000,
        burnin = 100000,
        thin = 100,
        pr = TRUE
)
```

## 2. M4-M6

```
prior4 <- list(
        B = list(mu = rep(0, 12), V = diag(12) * 1e+10), # Prior for fixed effects
        G = list(G1 = list(V = diag(2), nu = 2, alpha.mu=rep(0,2), alpha.V=diag(2)*1000)),
        R = list(V = diag(2), fix=2, nu = 2.002) # Prior for residual
)

prior5 <- list(
        B = list(mu = rep(0, 22), V = diag(22) * 1e+10), # Prior for fixed effects
        G = list(G1 = list(V = diag(2), nu = 2, alpha.mu=rep(0,2), alpha.V=diag(2)*1000)),
        R = list(V = diag(2), fix=2, nu = 2.002) # Prior for residual
)

prior6 <- list(
        B = list(mu = rep(0, 28), V = diag(28) * 1e+10), # Prior for fixed effects
        G = list(G1 = list(V = diag(2), nu = 2, alpha.mu=rep(0,2), alpha.V=diag(2)*1000)),
        R = list(V = diag(2), fix=2, nu = 2.002) # Prior for residual
)

formula4 <- cbind(log.lambda, is.variable) ~ trait - 1 + trait:Type + trait:median_length +
trait:FPKM_category + trait:RSA + trait:inter

formula5 <- cbind(log.lambda, is.variable) ~ trait - 1 + trait:Class + trait:median_length +
trait:FPKM_category + trait:RSA + trait:inter

formula6 <- cbind(log.lambda, is.variable) ~ trait - 1 + trait:Path + trait:median_length +
trait:FPKM_category + trait:RSA + trait:inter

#Example run
model <- MCMCglmm(
```

```
formula,
 random = ~us(trait):group,
rcov = ~us(trait):units,
family = c("gaussian", "threshold"),
prior = prior,
data = df_lambda,
nitt = 2100000,
burnin = 100000,
thin = 100,
pr = TRUE
)
```

## 3. M7

```
prior7 <- list(
    B = list(mu = rep(0, 30), V = diag(30) * 1e+10), # Prior for fixed effects
    G = list(G1 = list(V = diag(5), nu = 5, alpha.mu=rep(0,5), alpha.V=diag(5)*1000)),
    R = list(V = diag(5), fix=5, nu = 2.002) # Prior for residual
)
model7 <- MCMCglmm(
    cbind(log.dNdS, log.lambda, is.variable, log.prop, busted_positive) ~ trait - 1 +
    trait:Type + trait:median_length + trait:FPKM_category + trait:RSA + trait:inter,
    random = ~us(trait):group,
    rcov = ~us(trait):units,
    family = c("gaussian", "gaussian","threshold", "gaussian","threshold"),
    prior = prior,
    data = df,
    nitt = 2100000,
    burnin = 100000,
    thin = 100,
    pr = TRUE
)
```
